# Ab initio Spillover Compensation in CyTOF Data

**DOI:** 10.1101/2020.06.14.151225

**Authors:** Qi Miao, Fang Wang, Jinzhuang Dou, Ramiz Iqbal, Muharrem Muftuoglu, Rafet Basar, Li Li, Katy Rezvani, Ken Chen

## Abstract

Signal intensity measured in a mass cytometry (CyTOF) channel can often be affected by the neighboring channels due to technological limitations. Such signal artifacts are known as spillover effects and can substantially limit the accuracy of cell population clustering. Current approaches reduce these effects by using additional beads for normalization purposes known as single-stained controls. While effective in compensating for spillover effects, incorporating single-stained controls can be costly and require customized panel design. This is especially evident when executing large-scale immune profiling studies. We present a novel statistical method, named CytoSpill that independently quantifies and compensates the spillover effects in CyTOF data without requiring the use of single-stained controls. Our method utilizes knowledge-guided modeling and statistical techniques, such as finite mixture modeling and sequential quadratic programming, to achieve optimal error correction. We evaluated our method using five publicly available CyTOF datasets obtained from human peripheral blood mononuclear cells (PBMCs), C57BL/6J mouse bone marrow, healthy human bone marrow, chronic lymphocytic leukemia patient, and healthy human cord blood samples. In the PBMCs with known ground truth, our method achieved comparable results to experiments that incorporated single-stained controls. In datasets without ground-truth, our method not only reduced spillover on likely affected markers, but also led to the discovery of potentially novel subpopulations expressing functionally meaningful, cluster-specific markers. CytoSpill (developed in R) will greatly enhance the execution of large-scale cellular profiling of tumor immune microenvironment, development of novel immunotherapy, and the discovery of immune-specific biomarkers. The implementation of our method can be found at https://github.com/KChen-lab/CytoSpill.git.

## Introduction

The emergence and rapid adoption of mass cytometry (CyTOF) as a more scalable alternative to flow cytometry has led to unprecedented fine-grained profiling of human cell populations. CyTOF employs metal-isotope-tagged monoclonal antibodies to measure the expressions of the surface proteomic markers and/or intracellular signaling molecules in single cells. This technology is particularly important for various biomedical fields, such as immunology, oncology, and stem cell research, because CyTOF allows experimentalist to measure between 40 to 60 parameters (channels) from around 100,000 cells in a single assay. CyTOF has been widely used to profile the immune system in the presence of disease such as the characterization of cellular heterogeneity in tumor samples (Alizadeh et al., 2015; Wogsland et al., 2017). The advantage of this technology over its predecessor is the minimal amount of spectral overlap between channels that is more typical in flow cytometry and can lead to differing interpretations of cell populations (Kleinsteuber et al., 2016; Roederer, 2001). Despite this advantage, spillover effects similar to those in the flow cytometry data is still observed, since the intensity measured in a channel can be affected by the intensity of the neighboring channels.

Spillover effects, while generally minor, can substantially limit the accuracy of cell type identification. It is possible to alleviate this by selecting high purity isotopes, redesigning metal isotopes, or using control panel, but that approach is complicated, time-consuming, and costly (Chevrier et al., 2018). These approaches can be even more burdensome in the context of executing a large-scale immune profiling study where spillover effects might not be easily compensated for. Given that the spillovers have an approximately linear relationship with respect to the original signal (Chevrier et al., 2018), error reduction can be achieved by transforming the data using a properly estimated spillover matrix without necessitating any changes to study design or materials.

Here, we present a novel computational method, CytoSpill, that can independently compensate the spillover effects in CyTOF data without using any single-stained controls. Our method utilizes knowledge-guided modeling and statistical algorithms to infer the optimal spillover matrix and perform correction. We utilize the knowledge about spillover sources to constrain the estimation of the spillover matrix. The underlying assumption of our method is that the spillover component can be separated from the signal component in affected channels using a mixture distribution model.

## Materials and Methods

### Datasets

We examined our method using five CyTOF datasets, obtained from peripheral blood mononuclear cells (PBMCs) with a 36-antibody panel (Chevrier et al., 2018), C57BL/6J mouse bone marrow with a 38-antibody panel (Samusik, Good, Spitzer, Davis, & Nolan, 2016), healthy human bone marrow with a 32-antibody panel (Levine et al., 2015), chronic lymphocytic leukemia patient blood with a 46-antibody panel and healthy human cord blood sample with a 44-antibody panel. Table 1 presented the details of the datasets we used. All the datasets used were deposited in flow repository with ID: FR-FCM-Z2KW and on https://github.com/KChen-lab/CytoSpill.

**Table 1:** Descriptions of the datasets used in this study

| Datasets | Source | Number of Antibody Channels | Number of Cells | Tissue |
| --- | --- | --- | --- | --- |
| Peripheral blood mononuclear cells | Chevrier et al., 2018 | 36 | 87,817 | Human peripheral blood mononuclear cells |
| C57BL/6J mouse bone marrow | Samusik et al., 2016 | 38 | 86,864 | Mouse bone marrow |
| Healthy Human bone marrow | Levine et al., 2015 | 32 | 265,627 | Human bone marrow |
| Chronic lymphocytic leukemia patient blood | New | 46 | 126,855 | Human peripheral blood |
| Healthy human cord blood | New | 44 | 173,891 | Human cord blood |

### Spillover compensation problem

In CyTOF data, spillover usually comes from three sources: abundance sensitivity, isotopic impurity and oxidization (Takahashi et al., 2017). The information on these three sources can be obtained based on the isotopes used in the experiment panel. If an isotope used in one channel has a similar mass to the isotope used in another channel, for example differed by 1 atomic mass, these channels will affect each other (abundance sensitivity). Channels using isotopes with the same metal will affect each other (isotopic impurity). If the atomic mass of the isotope in a channel is larger than that in another channel by 16, the atomic mass of an oxygen atom, it will potentially be affected by this other channel through oxidation, since the metal may get oxidized incidentally during the experiment. In CyTOF experiments, cells expressing a marker will have positive readings while cells that do not express a marker will have zero readings. However, if a channel is affected by spillover, these cells may acquire some level of intensity readings, resulting from spillover effects. The signal artifacts resulting from spillover effects are usually smaller than the true biological signals, thus adding a low background modal to the marker expression density. We assume that the density of a marker affected by spillover follows a multimodal distribution, where the lowest intensity component corresponds to the spillover noise and the other components correspond to true expression levels.

The spillover compensation problem in CyTOF data is similar to the complete compensation problem defined in flow cytometry literate (Roederer, 2002). Let

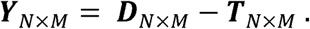

where ***D*** _*N* × *M*_ is the matrix of observed signals, ***T*** _*N* × *M*_ is the matrix of true signals and ***S*** _*N* × *M*_ is the spillover matrix whose diagonal elements are all 1’s. *N* is the number of cells and *M* is the number of channels (refer to Table 2 for commonly used notations in this paper). To model the spillover noise, we define the noise components as ***Y*** _*N* × *M*_ that

**Table 2:** Commonly used notations in this paper

| Notations | Definition |
| --- | --- |
| $\mathbf{D}$ | Matrix of observed signals |
| $\mathbf{T}$ | Matrix of true signals |
| $\mathbf{S}$ | Spillover matrix |
| $\mathbf{Y}$ | Noise component in the matrix of observed signals |
| $\mathbf{Z}$ | Channel interaction matrix |
| $\mathbf{I}$ | Identity matrix |
| $N$ | Number of cells |
| $M$ | Number of channels |
| $\ \cdot\ _F^2$ | Frobenius norm of a matrix |

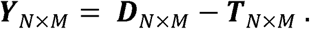

Thus, we have

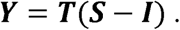

Since all diagonal elements of ***S*** are 1’s, (***S* − *I***)is a matrix contains only the off diagonal elements of ***S*** with ***I*** being an identity matrix. If we can estimate ***S***, we can perform compensation via ***T = DS***^−1^. ***S*** can be usually estimated by employing single-stained controls (Chevrier et al., 2018), which can be time-consuming and costly. Here, our goal is to derive ***S***through modelling of ***Y***. The mathematical problem is that only ***D*** is observed and that ***Y*** and ***T***, both having high dimensions, will need to be determined simultaneously from their relationship to ***D***. We hypothesize that it is possible to achieve this by making appropriate assumptions on ***Y*** and constraining the structure as well as the parameters in ***S*** using prior knowledge about the spillover structures. Based on our assumption that the noise component ***Y*** in each channel forms a lower modal in that channel’s intensity density distribution, we can model channel intensities using mixture probability distributions and segregate the noise modals from the mixed signals.

### Cutoff derivation

In order to separate the noise component from the true signal component, we derived a cutoff value for each channel in each CyTOF dataset. We assume that the intensity observed in each channel follows a mixture of Gaussian distributions where the Gaussian with the lowest mean represents the spillover noise. We fit the intensity distribution in a channel *j* (*j* = 1,2… *M*) by a finite mixture model using function initFlexmix from R package FlexMix, assuming there are *K* (*K* = 1,2… 5) components (Grün & Leisch, 2008). The observed marker expression level in channel *j* follows a multi-normal distribution:

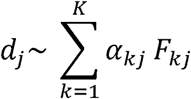

We choose the model with the lowest integrated completed likelihood value. If the model suggests more than one component (*K* > 1), we will derive a cutoff value *c* assuming a type 1 error *a* = 0.05.*K* = 2 suggests that the channel has a bimodal distribution and the lower modal is the noise component. The cutoff value *c* is then derived as the probability of *c* belonging to the lower modal:

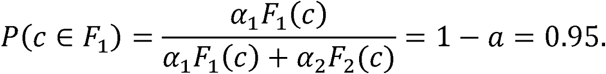

If *K* > 2, it suggests that the channel has a multimodal distribution. In that case, we will take the lowest modal and the highest modal to calculate *c*, assuming that the highest modal corresponds to the true signal. If the returned model only has one component, we will select an empirical cutoff at 10% quantile of the channel intensity. This is an important step in our method which defined the error components contributed by spillover effects on negative cells of the markers. These error components will be further utilized in next steps for estimating the spillover coefficients. Given the derived cutoff values ***C***= {*c*1, *c*2, *c*3 …. c*M*}, the noise component ***Y*** is defined as

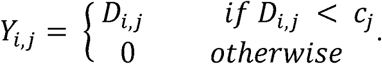

### Estimation of the Spillover Matrix

We attempted two methods to estimate the spillover matrix ***S***: 1) sequential quadratic programming with channel-specific constraints and 2) non-negative matrix factorization. The prior knowledge of error sources, i.e., abundance sensitivity, isotopic impurity and oxidization are utilized as constraints for model optimization.

#### Method 1: Sequential quadratic programming with channel-specific constraints

We define a channel interaction matrix ***Z*** _*M* × *M*_from the prior knowledge described above. *Z*_*i,j*_ is greater than 0 only when channel *i* can potentially affect channel *j* and 0 otherwise. Because spillover has strict additive effect (Chevrier et al., 2018), *Z* _*i,j*_ is not allowed to have negative values. Furthermore, previous studies indicated that the spillover effects are generally less than 10% (Chevrier et al., 2018). Thus, we apply a boundary constraint: *Z* _*i,j*_ ≤ 0.1 to the estimation. We estimate the channel interaction matrix using a sequential quadratic programming (SQP) algorithm (Kraft & Institut für Dynamik der Flugsysteme, 1988) as follows:

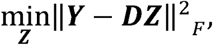

Where ***Z***satisfies 0 ≤ *Z*_*i,j*_ ≤ 0.1 when channel *i* can potentially affect channel *j* and *Z* _*i,j*_ = 0 otherwise. Here, ‖.‖^2^_*F*_denotes the Frobenius norm of a matrix. The spillover matrix ***S*** is calculated as ***S = Z* + *I***.

#### Method 2: Non-negative matrix factorization

Since ***D = TS*** where ***T*** and ***S*** are both non-negative by definition, we can also formulate the task of spillover matrix estimation as a masked non-negative matrix factorization (NMF) problem (Casalino, Del Buono, & Mencar, 2016; Lin & Boutros, 2020; Sobieraj, Kong, & Plumbley, 2017). We incorporate the cutoffs we derived and the prior knowledge into the non-negative matrix factorization model using binary masks. With the derived cutoff values ***C***, we generate a binary matrix ***B*** _*N* × *M*_ that masks non-noise component defined by the cutoff value in ***D*** that

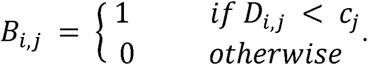

Given the channel interaction matrix ***Z*** defined above, we have:

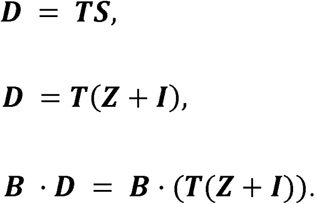

The noise mask ***B*** ensures that only the error component selected by our derived cutoffs can affect the optimization. The resulting method performs non-negative matrix factorization by solving the following optimization problem:

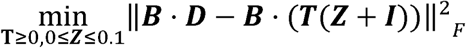

The optimization is performed using gradient decent algorithm in Tensorflow (Martín Abadi et al., 2015).

### Compensation

Given the estimated spillover matrix ***S***, we can obtain the real (compensated) data ***T*** via the following optimizing problem:

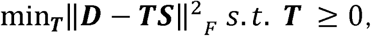

using a non-negative least squares (NNLS) algorithm (Chen & Plemmons, 2009; Chevrier et al., 2018). The ***T*** ≥ 0 since CyTOF data does not contain negative values.

### Evaluation

To evaluate our method, we compared the compensated data using the spillover matrix generated by our method with the compensated data using the single-stained controls. We also compared the uncompensated data with compensated data using four other datasets that do not have single-stained controls. We examined the data clustering results using t-SNE plots generated with Rtsne package followed by PhenoGraph clustering (Krijthe, 2014/2020; Levine et al., 2015; Maaten & Hinton, 2008).

### Simulation

We first used simulation to test the ability of SQP and NMF to estimate the spillover matrix in our model. We simulated true signals ***T*** with 20 channels and 100,000 cells. In each channel, the expression values of positive cells followed a normal distribution with mean randomly drawing from a uniform distribution U(100, 600) and standard deviation equals to mean/5. We assigned 0’s to the expression values of negative cells. For 10 of the 20 channels, we simulated them as channels with single modal positive expression. In each of these channels, we assigned 60,000 cells being positive cells and 40,000 cells being negative cells. For the other 10 channels, we simulated them with bimodal positive expressions where there are 30,000 cells in each modal followed by the normal distributions with different means. The remaining 40,000 cells were negative cells.

Based on our assumptions on error sources, we obtained the structure of the simulated spillover matrix. In each of the simulated spillover matrix ***S*** _*M* × *M*_ where *M* = 20, there are 116 off diagonal elements of spillover coefficients that need to be simulated. Moreover, the spillover coefficients were simulated based on a uniform distribution U(0, 0.1). We first simulated twenty spillover matrices. We then applied these spillover matrices to the simulated true signal ***T*** to generate the simulated CyTOF data ***D***_*simu*_ with spillover effects in it: ***D***_*simu*_ = ***TS***_*simu*_. We then performed estimation on ***S***_*simu*_ using our approaches and compared the estimated spillover matrices with the known simulated spillover matrices.

## Results

In order to compensate CyTOF data without using single-stained controls, we made assumptions that spillover noise contributes as a new modal at the lower end of the affected channel expression density. We derived a cutoff for each channel of the data to separate the noise modal from observed signal modal. We then assumed that the spillover effects come from three main sources which will constrain the spillover matrix structure that needs to be estimated. Both sequential quadratic programming with channel-specific constraints (cSQP) and non-negative matrix factorization (NMF) were applied to estimate the spillover matrix using the cutoff separated spillover noise component. Finally, the CyTOF data can be compensated using the obtained spillover matrix. Figure 1A shows the workflow of our compensation method and Figure 1B describes the assumptions of our method.

**Figure 1:**
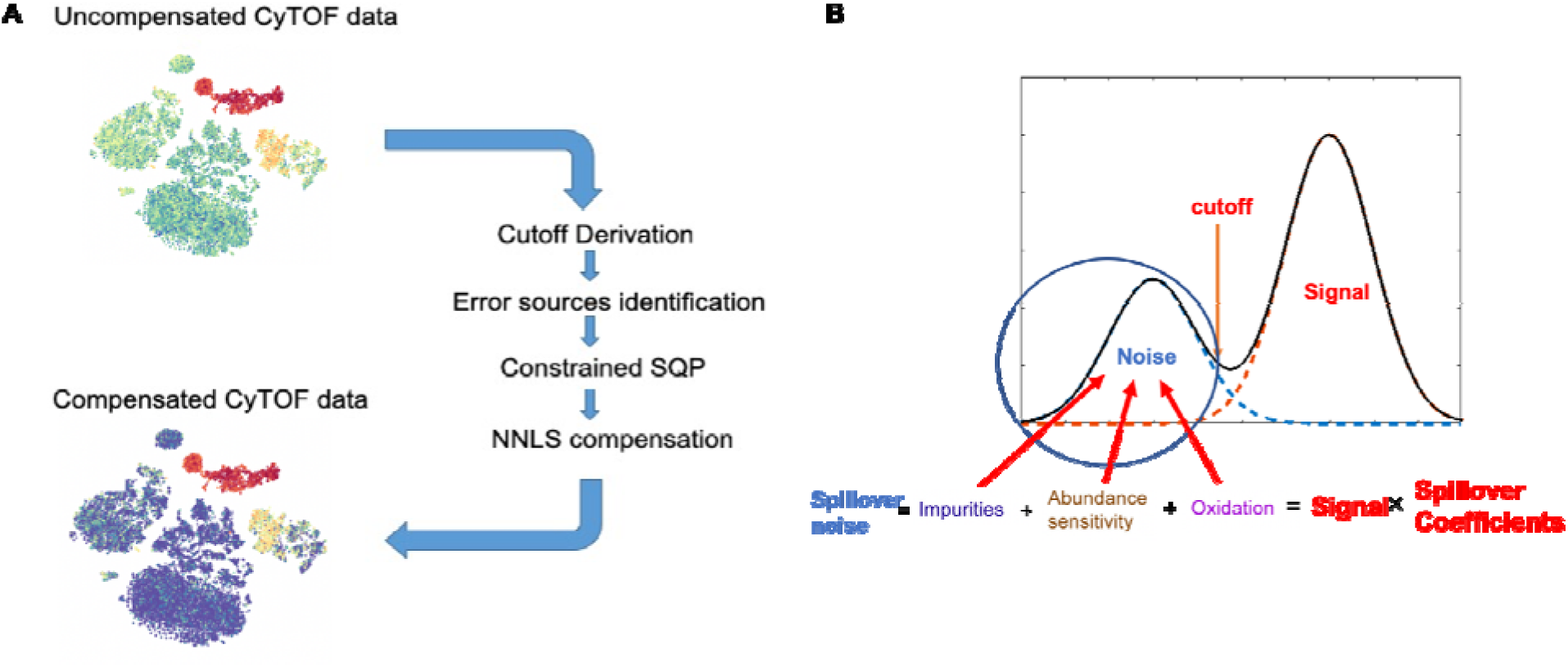
(A) Workflows of our compensation method. Based on our assumptions, we derived cutoffs based on uncompensated CyTOF data to identify the spillovers and estimate the spillover matrix using constrained sequential quadratic programming with prior knowledge. Non-negative least squares were used for compensation. (B) Assumptions of our methods. We assumed that a spillover affected channel will have a lower modal which contributed by spillover effects. We assumed the spillover were from three sources: isotope impurities, neighboring channel abundance sensitivity and oxidization.

### Simulation results

We applied two approaches, cSQP and NMF, to 20 different sets of simulated data. NMF did not reach final convergence, therefore, we took the results after 5,000 iterations for evaluation purposes. In our simulation analysis, we compared both cSQP and NMF estimated spillover matrix to the simulated ground truths. We found that cSQP estimated results were better aligned to the ground truth than NMF. From the scatter plots it can be seen that most data points aligned on the diagonal in cSQP results which suggest that cSQP estimated similar spillover matrix as the simulated ground truths while the majority of NMF estimated spillover coefficients lied on the boundary of 0 and 0.1 (Figure 2A, B). The cSQP approach achieved *R*^2^ = 0.51 compared to the NMF approach which achieved *R*^2^ = 0.02. Since the cSQP approach can estimate spillover coefficients much more accurately than the NMF approach, we chose the cSQP approach for our downstream assessments.

**Figure 2:**
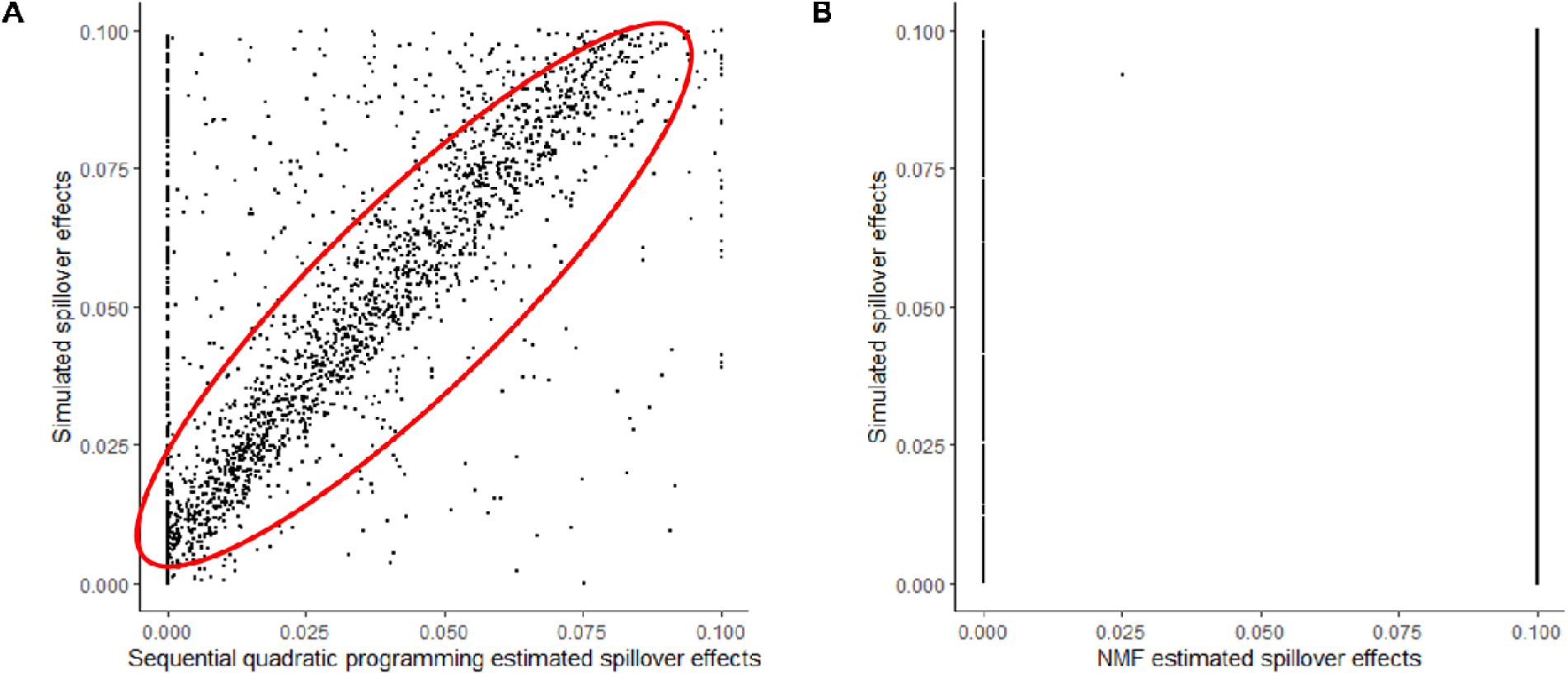
Comparison of simulated spillover effects and estimated spillover effects. axis represented the estimated spillover effects using our methods.axis represented the simulated true spillover effects. Each dot in the figure represented for an entry in a spillover matrix of our simulation study. (A) Scatter plot shows the result of sequential quadratic programming estimation and the= 0.51. (B) Scatter plot shows the result of non-negative matrix factorization and the= 0.02.

The spillover effect is a property of the metal labels and the instrument and does not depend on biological markers or samples. We further demonstrated that the spillover effect is not affected by relative abundance of markers using simulation. We simulated data with different level of signals and obtained spillover effects that are independent of the level of signals (Fig S1). The detailed description of the simulation can be found in Supplementary Materials 1.

### Comparison with single-stained controls dependent compensation

We analyzed a PBMCs dataset stained using a 36-antibody panel with single-stained controls to compare the accuracy of the spillover matrices obtained using CytoSpill with those obtained using single-stained controls. In Figure 3A, we show the t-SNE plot of this data before compensation and it has 20 PhenoGraph clusters. The panel used for this data was developed for immune cell type identification, and some of the proteins on this panel were conjugated with two different metal labels. In Figure 3B, we can observe the spillover intuitively by comparing the expression of the two metal labels on the same protein using expressing plots. The different expression between the two metals used for CD8, CD3 and HLA-DR suggested that there are spillovers in 174Yb-conjugated CD8, 173Yb-conjugated CD3 and 171Yb-conjugated HLA-DR. After compensation performed by CATALYST using single-stained controls, the same protein conjugated with different metals has almost identical expression profiles by comparing the expression plot. After compensation performed using CytoSpill generated spillover matrix, our method also removed the spillover effects and achieved similar results as using single-stained controls without requiring their use.

**Figure 3:**
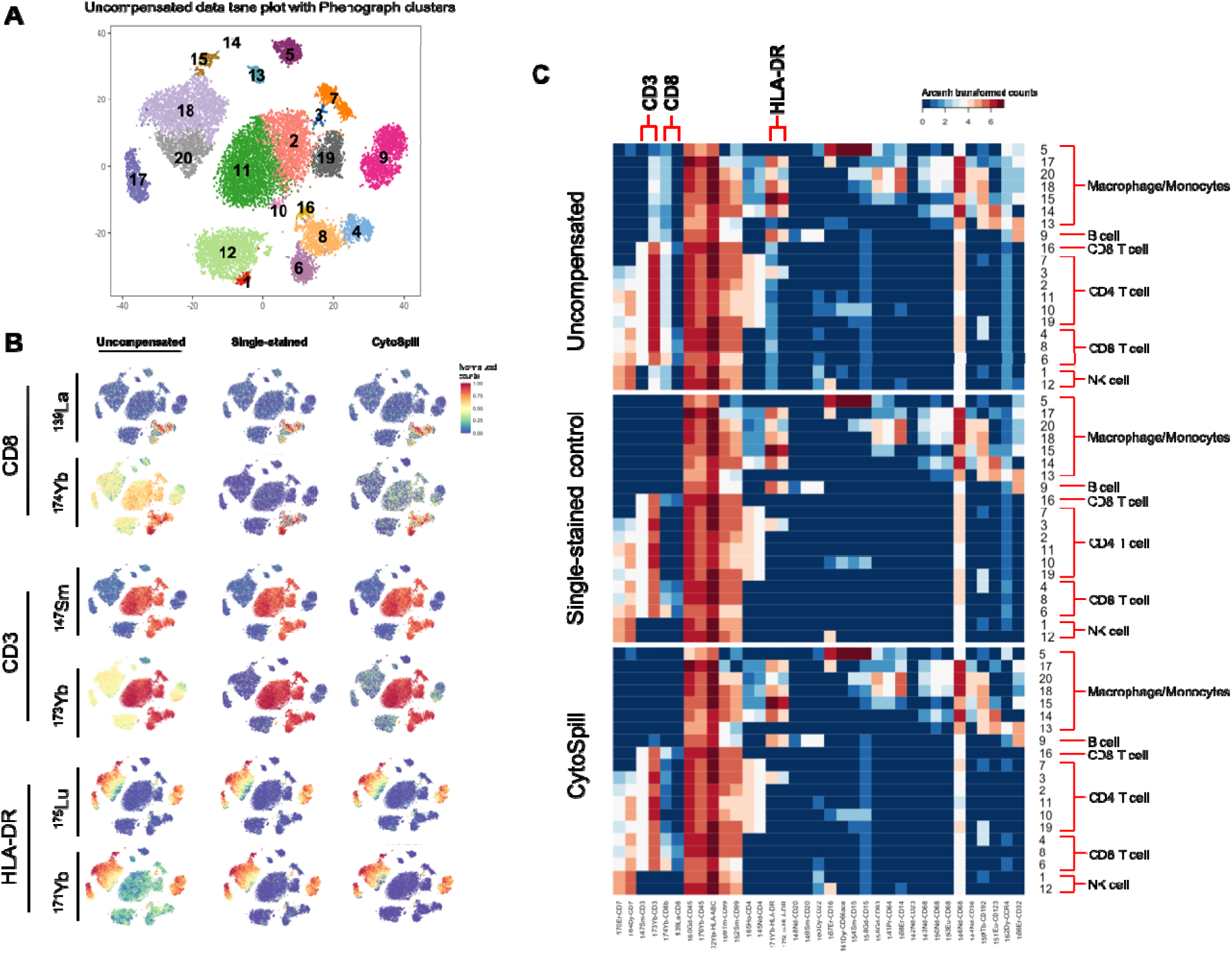
Comparison of CytoSpill generated spillover matrix compensated data with single-stained controls generated spillover matrix compensated data. (A) t-SNE plot of uncompensated data labeled with PhenoGraph clusters. (B) Marker expressions based on t-SNE projection by uncompensated data. Expression values were normalized between 0 and 1. Spillovers were observed in 174Yb-stained CD8, 173Yb-stained CD3 and 171Yb-stained HLA-DR. CytoSpill achieved comparable results with single-stained controls on compensate these markers. (C) Heatmaps showed compensation results on PhenoGraph clusters based on uncompensated data. Expression values were arcsinh transformed. Clusters were annotated with cell types.

In Figure 3C, the heatmap of marker expressions in different clusters of uncompensated, single-stained controls compensated and CytoSpill compensated data can be seen. We annotated the clusters with cell types based on the marker expressions. We found that 173Yb-conjugated CD3 has expression in non-T cells clusters compared to 147Sm-conjugated CD3. The 174Yb-conjugated CD8 marker were observed to have expression in the non-CD8 T cells clusters compared to 139La-conjugated CD8. The 171Yb-conjugated HLA-DR had expressions in clusters besides the Macrophage/Monocytes and B cells clusters, while 175Lu-conjugated HLA-DR did not. The expression pattern in these clusters should be negative but were both removed using single-stained controls in CATALYST and ab initio in CytoSpill. This confirms that CytoSpill can effectively remove the true spillover in the CyTOF data without using single-stained controls.

### Applying CytoSpill on four immune datasets

Moreover, we also applied CytoSpill on four additional immune related datasets, including samples from human bone marrow, mouse bone marrow, human peripheral blood and human cord blood. We checked the marker expression before and after compensation. Also, t-SNE plots were generated and PhenoGraph was applied for clustering analysis (Krijthe, 2014/2020; Levine et al., 2015; Maaten & Hinton, 2008). We observed that some markers in these datasets have a high expression in certain clusters, while they also have some amount of intermediate expressions in other clusters. These intermediate expressions were lowered or removed after running CytoSpill.

For example, the CD8 markers in the leukemia patient peripheral blood data and the healthy human cord blood data were strongly affected by spillover (Figure 3, 4). Figure 3A shows the t-SNE plot of uncompensated leukemia patient peripheral blood data, which is labeled by manually gated cell populations. We found that CD8 marker has an intermediate level in the B cells and CD4 T cells clusters, which may be caused by spillover. After performing CytoSpill compensation, these spillover signals were removed (Figure 3B, 3D). Figure 3C shows the histogram of the uncompensated and compensated arcsinh transformed CD8 expression with the dotted line representing the derived cutoff value for this channel. The noise component below the cutoff was substantially lowered after compensation with more cells having a level close to 0 and the positive population above the cutoff remained unchanged.

**Figure 4:**
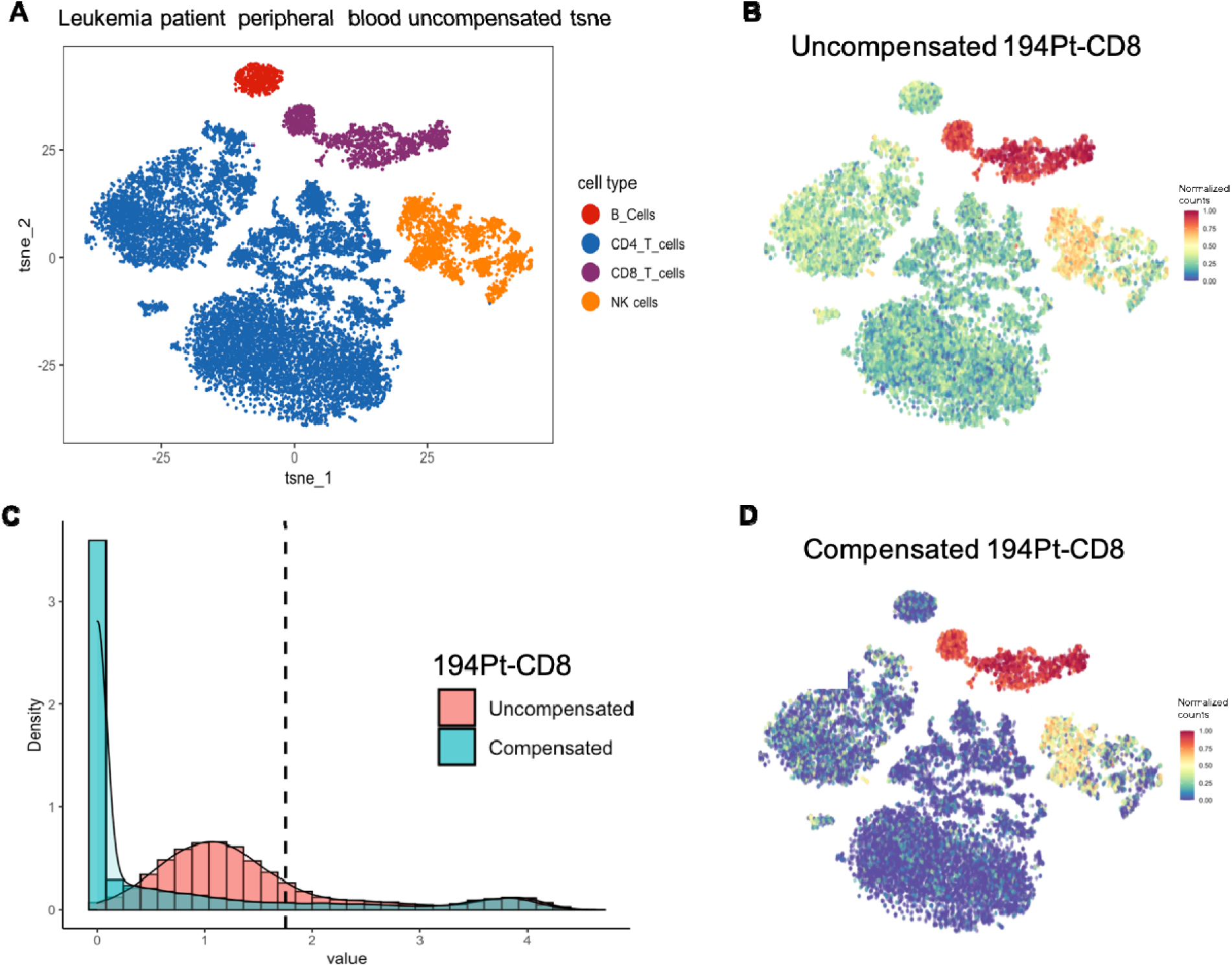
CD8 marker spillover in leukemia patient blood data were compensated using CytoSpill. (A) The t-SNE plot generated with uncompensated data and labeled by manual gated populations. (B) The normalized expression plot of 194Pt-conjugated CD8 marker before compensation. Spillover were observed outside NK cells and CD8 T cells. (C) Histograms showed the CD8 marker expression density before and after compensation. Dotted line represents the derived cutoff value. The noise component below the cutoff were lowered after compensation. (D) The normalized expression plot of 194Pt-conjugated CD8 marker after compensation. We observed that spillover were lowered after compensation.

**Figure 5:**
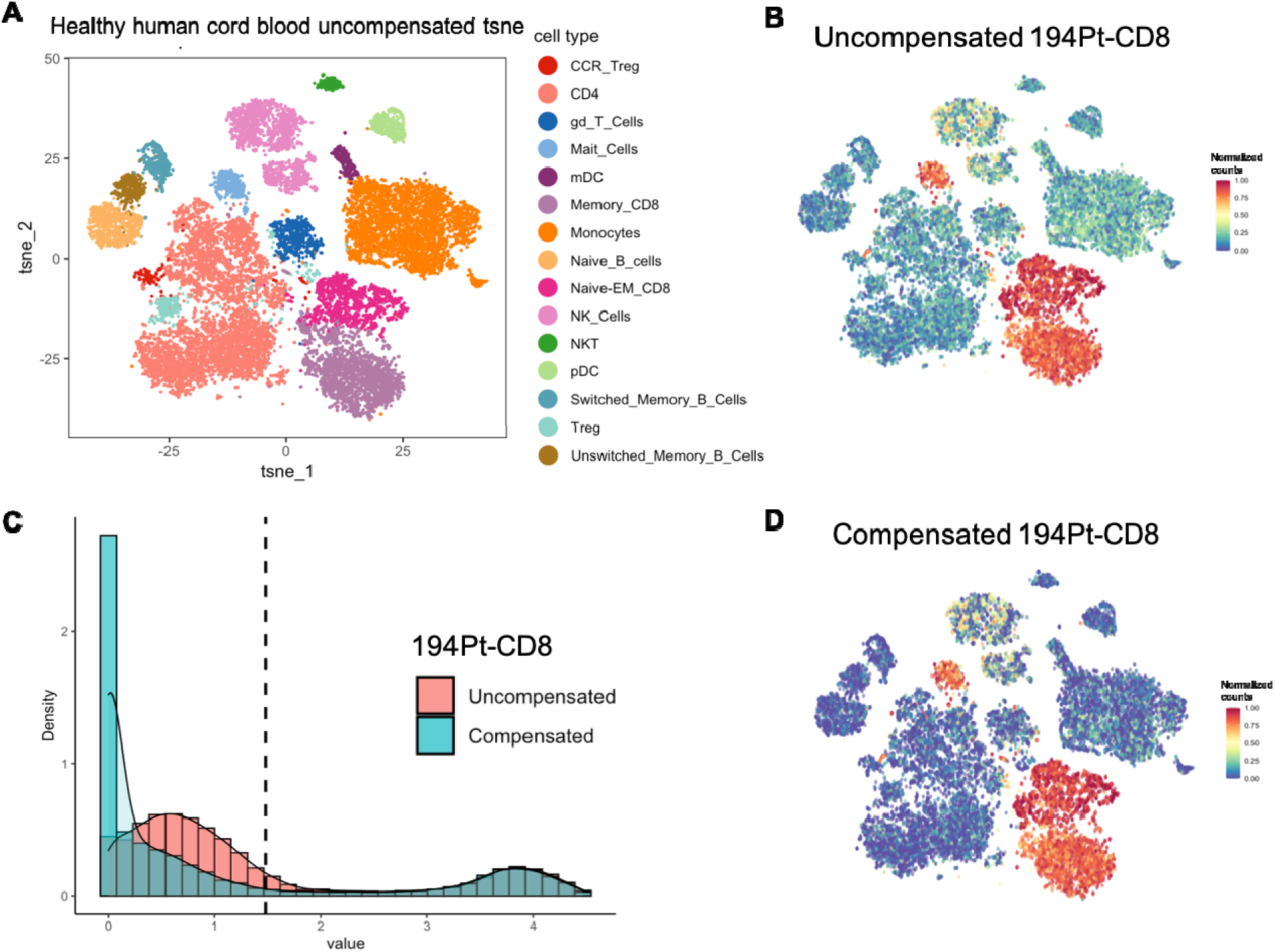
CD8 marker spillover in healthy human cord blood data were compensated using CytoSpill. (A) The t-SNE plot generated with uncompensated data and labeled by manual gated populations. (B) The normalized expression plot of 194Pt-conjugated CD8 marker before compensation. Spillover were observed outside CD8 T cells, NK cells and Mait cells. (C) Histograms showed the CD8 marker expression density before and after compensation. Dotted line represents the derived cutoff value. The noise component below the cutoff were lowered after compensation. (D) The normalized expression plot of 194Pt-conjugated CD8 marker after compensation. We observed that the spillover were lowered after compensation.

In the healthy human cord blood data, the CD8 marker has a high expression on the mucosal associated invariant T cell and the CD8 T cell clusters (Figure 4A, 4B). CD8 also expressed on some of the NK cells. However, we found it also has intermediate levels on other clusters which should be negative for the CD8 marker. After performing compensation, the spillover caused signals were removed on these clusters (Figure 4B, 4D). Figure 4C showed the histogram compared the CD8 expression density before and after compensation. Noise component below the derived cutoff were lowered after compensation and the positive population above the cutoff remained.

### Discovering novel clusters

In terms of PhenoGraph clustering results, we found that the PhenoGraph clusters number increased from 22 to 23 after performing compensation on the healthy human bone marrow data, from 20 to 21 on the mouse bone marrow data and from 26 to 29 on the leukemia patient peripheral blood data. The increased number of clusters suggests that performing compensation may lead to discovery of novel cell clusters. In the healthy human bone marrow data, we found that two new clusters of T cells were revealed after compensation (Figure 6). Figure 6A-D shows the t-SNE plots for the clustering results obtained before and after compensation. Based on the t-SNE plots and PhenoGraph clustering results, two new clusters 2 and 4 from CD4 T cells and CD8 T cells respectively were formed after compensation (Figure 6C, 6D). By checking the expression signature of these two clusters we found that they are T cells with low CD44 expression which were not revealed as clusters before compensation (Figure 6E, 6F). CD44 is an activation marker which distinguishes naïve T cells from memory and effector T cells (Baaten, Li, & Bradley, 2010; Schumann, Stanko, Schliesser, Appelt, & Sawitzki, 2015). These two novel clusters we found are thus likely naïve T cells clusters, which were masked by the spillover effects in the uncompensated data.

**Figure 6:**
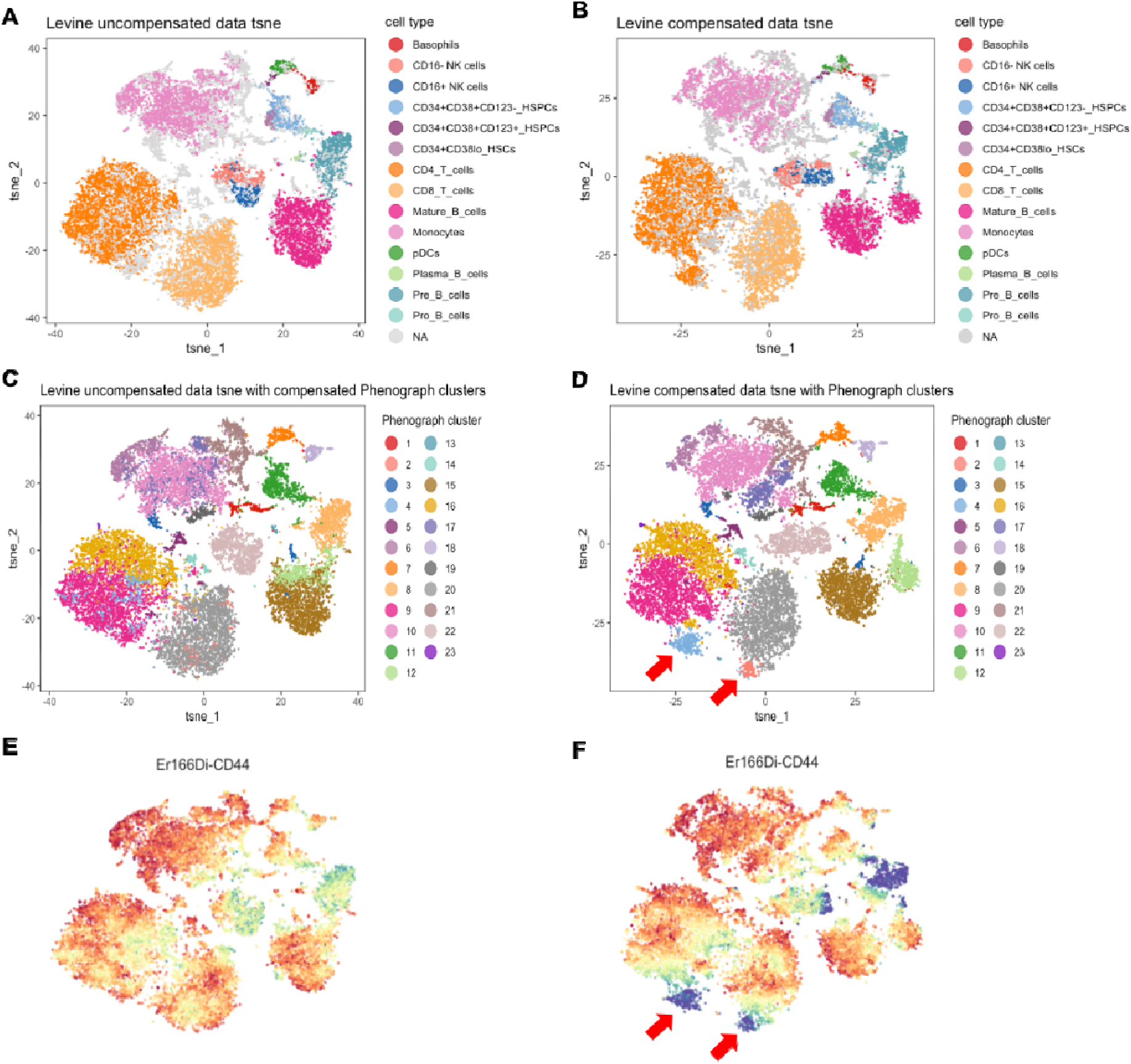
Novel clusters discovered after compensation in healthy human bone marrow data. (A, B) showed the t-SNE plots generated based on uncompensated and compensated data labeled by cell populations. (C, D) showed the t-SNE plots labeled by compensated data generated PhenoGraph clusters. Two new clusters 2 and 4 were found after compensation. These two clusters of cells were originally scattered in CD4 T cells and CD8 T cells before compensation. (E, F) showed the CD44 expression level before and after compensation. The newly found clusters have lower CD44 expression.

Identifying novel, meaningful clusters after performing compensation indicate that our compensation method can led to more precise cell-type clustering, in conjunction with the application of the t-SNE and the PhenoGraph algorithms.

## Discussion

In immunology research, CyTOF was widely used to dissect the heterogeneity of immune cell populations. It is well accepted that the spillover effects in CyTOF data could led to inaccurate population dissection. In this paper, we presented a new method that could alleviate spillover effects in CyTOF data without relying on additional control data such as single-stained controls. In our method, we used finite mixture model to derive the cutoffs that separates the noise and signal for each channel. We then use a constrained sequential quadratic programming approach to infer the spillover matrix that optimally quantifies the spillover effects. We performed simulation studies to demonstrate the ability of our method to deconvolve spillover effects on multiple published datasets and unpublished datasets. We observed markers affected by spillover effects and removed the noise after performing compensation. The compensation also led to increased number of PhenoGraph clusters in multiple datasets. In the healthy human bone marrow, our method discovered novel and meaningful cell subpopulations that would have been buried in the uncompensated data. To our knowledge this is the first method that can compensate spillover effects in CyTOF data without requiring single-stained controls.

We have considered two different approaches, constrained sequential quadratic programming (cSQP) and masked NMF, to integrate prior knowledge into our model and derive the spillover matrix. In our simulation study, we demonstrated that the cSQP approach had better performance than the NMF approach. NMF was not able to converge under our constraints. Since our spillover matrix is sparse under our assumption, some kind of sparsity constraint is required to be implemented in the optimization of NMF. There are several prior studies implement NMF based method on spectral unmixing, these studies have used sparsity constraints (Jiménez-Sánchez, Ariz, Morgado, Cortés-Domínguez, & Ortiz-de-Solórzano, 2020; Lu, Wu, & Yuan, 2014; Rajabi & Ghassemian, 2015). However, the spillover matrix structure was pre-defined based on prior knowledge in our study. To implement these prior knowledges in NMF, we used masked NMF (Casalino et al., 2016; Lin & Boutros, 2020; Sobieraj et al., 2017). Our constraint on the spillover matrix was too strict, leading to the undesirable performance of the masked NMF in our study.

Our method assumed that the spillover effects are caused by three main sources that are considered to be relatively mild (being a small fraction of the observed signals). Besides spillover effects, the noise in CyTOF data could also come from contaminations from other samples in the lab or external environment (Leipold, Newell, & Maecker, 2015; Takahashi et al., 2017). However, these could be controlled based on rigorous experimental protocols (Takahashi et al., 2017). Our method assumed the noise in the data come primarily from spillover. On datasets where these assumptions are violated (e.g., errors constitute a large fraction of the observed signal), the performance of our method may be impaired.

Our assumptions on spillover sources were based specifically on the CyTOF technology, which is different from the flow cytometry technology. Our method also assumed non-negativity of CyTOF data, while data from flow cytometry could have negative values due to background subtraction (Tung et al., 2007). Thus, our method is effective for CyTOF data analysis and will not be applicable to flow cytometry data.

For the future work of our method, we would like to explore whether our proposed compensation method can lead to improvement in predicting clinical outcome and discovering novel disease mechanisms.

CytoSpill is implemented in R and the source code is released on GitHub: https://github.com/KChen-lab/CytoSpill. We expect that our method will significantly benefit the cancer and immunology research community in studying tumor microenvironment and developing novel immunotherapy.

## Acknowledgements

This work was supported in part by the University Cancer Foundation via the Institutional Research Grant program at the University of Texas MD Anderson Cancer Center, Human Breast Cell Atlas Seed Network Grants (CZF2019-002432 and CZF2019-02425) from the Chan Zuckerberg Initiative DAF, an advised fund of Sillicon Valley Community Foundation, grant RP180248 to KC from Cancer Prevention & Research Institute of Texas, a fellowship on the NLM Training Program T15LM007093 to RI, and P30 CA016672 (US National Institutes of Health/National Cancer Institute) to the University of Texas Anderson Cancer Center Core Support Grant.

## Supplementary Materials

### 1. Simulation to demonstrate whether levels of expression will affect the spillover effects estimation

To demonstrate that the spillover effects are not affected by relative abundance of markers, we performed a simulation with a 4-channel panel and 20,000 cells. One of the four channels was affected with spillover by the three other channels.

Total four different true signal settings were simulated. We simulated the affected channel with three different levels of expression that 10,000 positive cells followed distributions of N(150, 75), N(300,75) and N(600,75) respectively and 10,000 negative cells had expression values of 0. We also left the affected channel empty with expression value of 0 on all cells.

The true signals of the other three channels were simulated that in each channel, 10,000 positive cells followed distributions of N(200, 75), N(300, 75), N(400, 100) and 10,000 negative cells had expression values of 0. The negative cells in the affected channel will have positive expressions on the other three channels.

We simulated the true signal and 10 spillover matrices such that we could generate an approximation of real-world data. We then compared the simulated spillover matrices against the estimated spillover matrices using our method based on the different expression levels of the affected channels.

In Figure S1, the comparison between estimated spillover effects and simulated spillover effects under different levels of expression can be seen. A diagonal line means the estimated spillover effects match well with the simulated spillover effects. In all four conditions, our method performed well in the spillover effects estimation and have *R*^2^ equals to 0.99. This simulation study suggests that the different level of expressions (or no expression at all) will not affect the estimation of spillover effects in our method.

**Figure S1:**
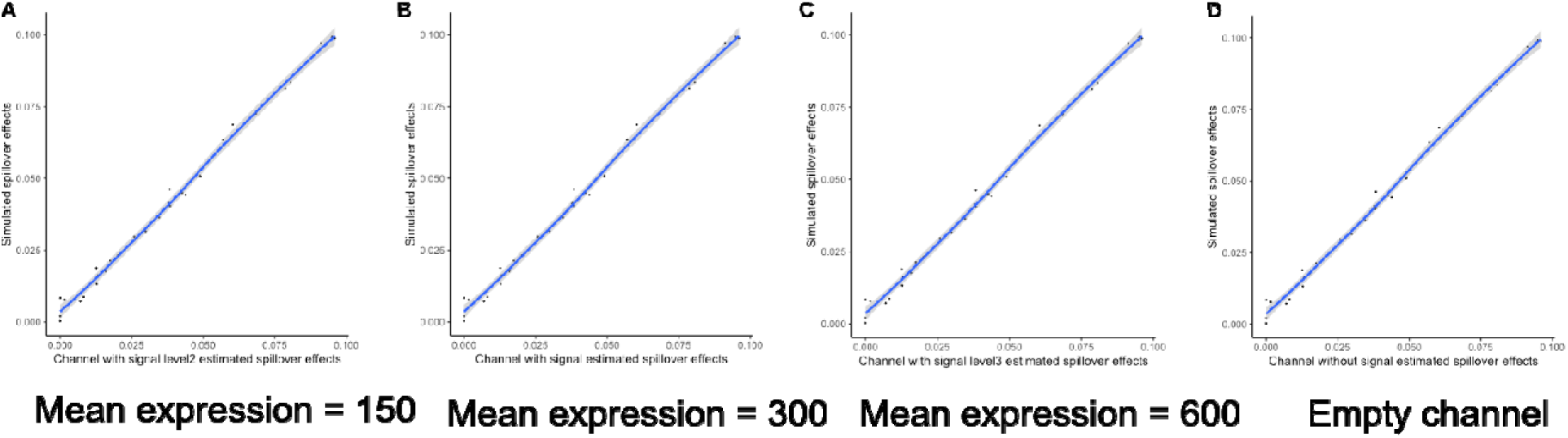
Comparison of simulated spillover effects and estimated spillover effects. axis represented the estimated spillover effects using our methods.axis represented the simulated true spillover effects. (A-D) Plots showed the comparison of estimated spillover effects compared with simulated spillover effects under different levels of expression of the affected channel. All of the results have which suggest the estimation performed well.

## Notes

1 This research was supported by the University Cancer Foundation via the Institutional Research Grant program at the University of Texas MD Anderson Cancer Center, Human Breast Cell Atlas Seed Network Grants (CZF2019-002432 and CZF2019-02425) from the Chan Zuckerberg Initiative DAF, an advised fund of Sillicon Valley Community Foundation, grant RP180248 to KC from Cancer Prevention & Research Institute of Texas, a fellowship on the NLM Training Program T15LM007093 to RI, and the Cancer Center Support Grant P30 CA016672 to PP from the National Cancer Institute.

### Competing Interest Statement

The authors have declared no competing interest.

